# Calibration, Selection and Identifiability Analysis of a Mathematical Model of the *in vitro* Erythropoiesis in Normal and Perturbed Contexts

**DOI:** 10.1101/314955

**Authors:** Ronan Duchesne, Anissa Guillemin, Fabien Crauste, Olivier Gandrillon

## Abstract

The *in vivo* erythropoiesis, which is the generation of mature red blood cells in the bone marrow of whole organisms, has been described by a variety of mathematical models in the past decades. However, the *in vitro* erythropoiesis, which produces red blood cells in cultures, has received much less attention from the modelling community. In this paper, we propose the first mathematical model of *in vitro* erythropoiesis. We start by formulating different models and select the best one at fitting experimental data of *in vitro* erythropoietic differentiation. It is based on a set of linear ODE, describing 3 hypothetical populations of cells at different stages of differentiation. We then compute confidence intervals for all of its parameters estimates, and conclude that our model is fully identifiable. Finally, we use this model to compute the effect of a chemical drug called Rapamycin, which affects all states of differentiation in the culture, and relate these effects to specific parameter variations. We provide the first model for the kinetics of *in vitro* cellular differentiation which is proven to be identifiable. It will serve as a basis for a model which will better account for the variability which is inherent to experimental protocol used for the model calibration.

## 1 Introduction

Erythropoiesis is the process by which red blood cells are produced. It occurs within the broader frame of haematopoiesis, the process which generates all blood cells. The dynamics of haematopoiesis has been extensively modelled mathematically in the past decades, with the first historical models published as early as fifty years ago^(1,2)^ (for a review of the history of haematopoiesis modelling in general, see^(3)^). Models of haematopoiesis have improved the understanding of both the processes they describe^(4)^, and the mathematical tools they use. These models comprise non-exhaustively Differential Equations (DE, either ordinary^(1,2,5)^, partial, which can be structured by age, maturity, or a combination of these^(6)^, or even delay differential equations^(7)^) and agent-based models^(8)^. They can be fully deterministic^(5)^ or can include a more or less prominent stochastic component^(9,10,11)^.

Recent works specifically focusing on erythropoiesis comprise DE-based models^(12,13)^ and multi-scale descriptions of these phenomena^(14,15,16)^.

All these works aim at modeling the *in vivo* physiological processes, *i.e.* the processes occurring in a whole organism. Those processes are related to numerous pathologies, for which clinical data are very sparse and must be acquired on experimentally prohibitive time-scales, which complicates their study. Modeling has therefore provided significant insights into these pathologies^(4)^. On the contrary, the *in vitro* context, *i.e.* the process that takes place in cells grown in culture, is much simpler to characterize experimentally. Yet, to our knowledge, no modeling study has focused on it so far. Since the *in vitro* differentiation is an experimental tool of choice for the study of cellular decision-making^(17,18,19)^, we propose to develop a model for the dynamics of the *in vitro* erythropoiesis.

Moreover, the current models of erythropoiesis suffer from one major drawback: the weakness of their parameterization, which can fall within three categories.

A vast majority of the existing models of erythropoiesis are based on experimental parameter values from the litterature. In some cases these values are used in other contexts that those in which they were obtained (typically, in other species^(12)^).

In other cases, the parameter values of a model are chosen arbitrarily to reproduce a qualitative behaviour. Apart from this qualitative fit, such approaches do not provide any information regarding the validity of the values^(16)^.

Finally, when the parameters of a model are estimated to reproduce a dataset, the precision of this estimation is seldom investigated^(20)^. By this, we mean that depending on the algorithmic details of the estimation, it is possible that several values of the parameters might render the same fit to the data. In this case the model is said to be unidentifiable.

A model is said to be identifiable if and only if it is possible to infer a unique value for each of its parameter by comparing its output to experimental data. Otherwise it is unidentifiable. A model can be non-identifiable for several reasons^(21,22)^.

Structural identifiability is related to the structure of the model, and the observed variables. A model is structurally unidentifiable when several of its parameters are redundant, meaning that they can vary in such a manner that the measured output of the model is not affected^(21,22,23)^. A variety of methods, based on different approaches, can be used to assess the structural identifiability of a dynamic model. These include, non-exhaustively, the Taylor series method^(24)^, the similarity transformation^(25)^, the generating series method^(26)^, and the profile likelihood approach^(22,27,28)^. A review of these methods is provided in^(21)^ and their performance is assessed in^(23)^.

Practical Identifiability is related to the quantity and quality of the data used for model calibration. If the data is too sparse or too noisy to estimate all parameters together, then the model is said to be practically unidentifiable^(21,22)^. Essentially three kinds of frequentist methods can be used to assess the practical identifiability of a model: methods based on the Fisher Information Matrix (FIM)^(29)^, which use a parabolic approximation of the likelihood function, profile-likelihood-based methods^(22,28)^, and bootstrapping, which is based on the resampling of the data^(30)^. FIM-based methods are less computationally demanding, because they require the fewest parameter estimation steps^(31)^, but due to their parabolic approximation they are proven to render biased results^(22)^. On the other hand, the profile likelihood-based method is computationally cheaper than bootstrapping^(31)^, and is proven to detect both structural and practical unidentifiabilities^(22,27)^.

Despite the growth of the interest in identifiability and related concepts among the biological systems modelling community^(21,32,33,34)^ the identifiability of models remains seldom investigated^(20)^. With this in mind, the most rigorous way to design and calibrate a model of erythropoiesis seems to use dedicated experiments to determine its parameter values. Once these values have been determined, one should then test the identifiability of the model before using it for any prediction.

In this paper, we aim at developing an identifiable model for the dynamics of the *in vitro* erythroid differentiation. The data that we use to calibrate it consists in counts of different cell sub-populations at regularly spaced time-points during the course of proliferation and differentiation of chicken erythroid progenitors. We start by formulating different possible structures for the dynamics of the system, and for the distribution of residuals. We select the best structure and distribution using classical information criteria. We then assess the identifiability of our model using an approach based on the profile likelihood concept. Finally, we test the adaptation of our model in a perturbed context, when cells are exposed to rapamycin, a drug which is known to affect the dynamics of differentiation, although its precise effect on proliferation and differentiation remains unclear. Since our model is identifiable, it is possible to quantify the effect of the drug on each of its parameters.

## 2 Methods

### 2.1 Data

The experimental setting from which were obtained all the data used in this study consists in a population of chicken erythroid progenitors which may either be maintained in a proliferative state or induced to differentiate into mature erythrocytes. Their behavior is controlled by the medium in which they are grown^(35)^. In the self-renewal medium (referred to as the LM1 medium) the progenitors keep undifferentiated, and undergo division after division. The LM1 medium contains growth factors (TGF-*α* and TGF-*β*), which stimulate the proliferative ability of the progenitors. On the contrary, the differentiation medium (referred to as the DM17 medium), contains anaemic chicken serum and insulin. After switching medium from LM1 to DM17, a fraction of the progenitors undergoes differentiation and becomes erythrocytes. The culture thus becomes a mixture of differentiated and undifferentiated cells, with some keeping proliferating. These two media can be used in different cellular kinetics experiments.

The first one, referred to as the LM1 experiment, consists in culturing cells in their self-renewing medium. Trypan blue is a non-toxic chemical that specifically dyes dead cells (fig 1-A). Dying a sample of the culture and counting the stained cells under a microscope thus gives access to the absolute numbers of living cells, at different times during their proliferation (namely, one point per day from day 0 to day 4 after the beginning of the experiment).

The second one will be referred to as the DM17 experiment, and is based on the culture of progenitors in their differentiation medium. In this experiment, the culture is a mixture of dead and living, self-renewing and differentiating cell. A chemical called benzidine, which specifically dyes the differentiated cells, gives access to the number of differentiated cells in the medium (fig 1B). A parallel staining with trypan blue still gives access to the overall numbers of living cells (fig 1B). Consequently, the data available from this experiment are the absolute numbers of differentiated cells, as well as the total number of living cells (which comprises both self-renewing and differentiated cells) at the same time points as in the LM1 experiment.

These experiments have been performed in two settings. In the first one cells are grown in their regular media (Figure 1C, black dots). In the other one, they grow in the presence of rapamycin, a chemical drug known to affect both the number of living cells in culture, and the proportion of differentiated cells^(36,37)^, as displayed on Figure 1C.

**Figure 1:**
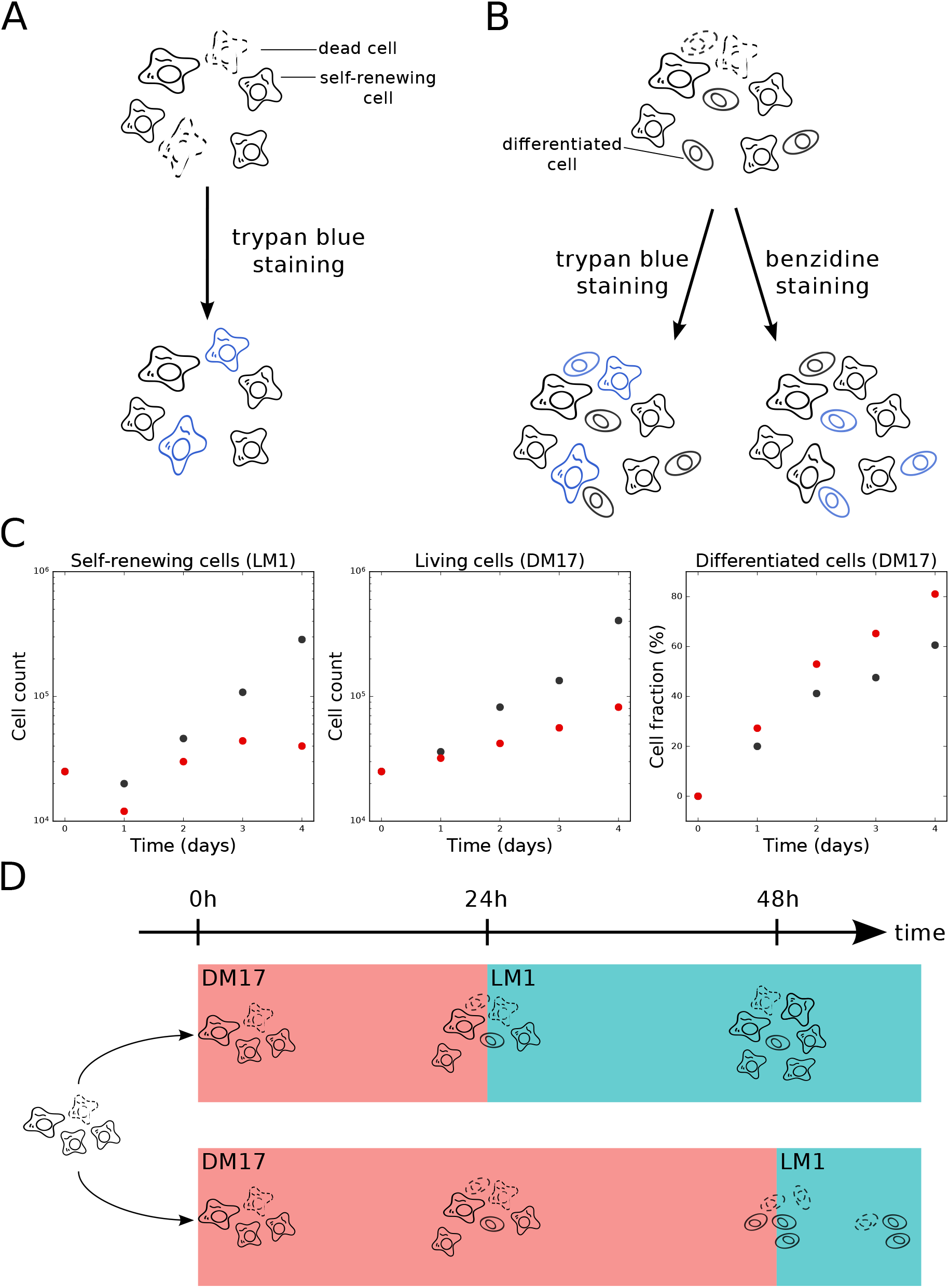
Experimental context. A: In LM1 medium the culture is only composed of living and dead cells in the self-renewal state. The amount of living cells can be measured by trypan blue staining. B: In DM17 medium, the culture is a mixture of living and dead, self-renewing and differentiating cells. The amount of living cells can be measured by trypan blue staining. The amount of differentiated cells can be measured by benzidine staining. C: Data used to calibrate the models. Black dots are the results of the cell counting experiments in the control situation (no treatment). Red dots are the results of the same experiments under rapamycin treatment. Both conditions were obtained with the same initial populations, so the black and red dots are the same at *t* = 0. For readability, living cell counts are displayed in log-scale, and differentiated cell counts are displayed as a fraction of the total living cell count. D: Commitment experiment. If the differentiating cells are switched back to LM1 after 24h of differentiation the culture starts proliferating again (upper trajectory). If the cells are switched back to LM1 after 48h, the culture stagnates (lower trajectory).

Another piece of experimental data that will be of use for calibrating our models is the result of the commitment experiment (Figure 1D). In this experiment, once a cell culture has been switched to the DM17 medium, it can be switched back to the LM1 medium^(17)^. Switching back after 24 hours of differentiation does not cancel the proliferative ability of the progenitors (the culture gets proliferating again), but switching back after 48 hours does (instead of proliferating again, the culture stagnates), as summarized on Figure 1D. This means that there must remain some self-renewing cells in the culture after one day, but that they all have started differentiating after two days.

### 2.2 Models

#### 2.2.1 Structural Model

We propose three alternative dynamic models of the erythroid differentiation, which are summarized on Figure 2.

**Figure 2:**
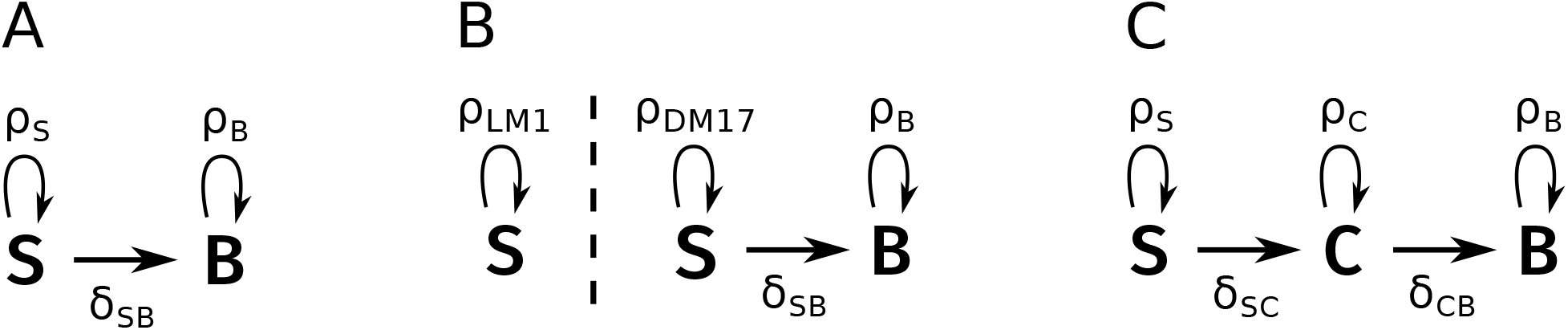
Diagrams of three possible structural models for our data. A: The SB model has no intermediary compartment. B: The S2B model has no intermediary compartment, but the self-renewing cells change proliferation rate in DM17. C: The SCB model has an intermediary compartment.

The SB model comprises only two compartments (Figure 2A), a self-renewing one (S) and a differentiated one (B, which stands for *Benzidine-positive*), whose dynamics are given by the equations:

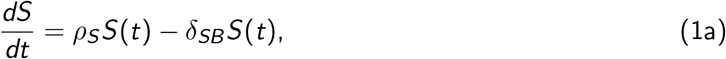

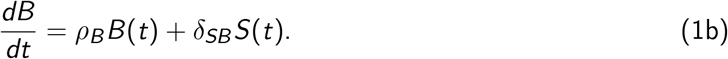

This model is characterized by a set *θ* = (*ρ_S_*, *δ_SB_*, *ρ_B_*) of three parameters, where *ρ_i_* is the net proliferation rate of compartment *i*. For estimation-related reasons, it incorporates the balance between cell proliferation and cell death. This means that *ρ_i_* can be either positive (more proliferation than death) or negative (more death than proliferation). On the other hand *δ_ij_* is the differentiation rate of cell type *i* into cell type *j*, which is positive.

The S2B model comprises also the S and B compartments (Figure 2B), but allows the self-renewing cells to change their net proliferation rate upon culture medium switching. This formulation arose from the consideration that proliferation is faster in the DM17 than in the LM1 medium (Figure 1C). The dynamics of this model are given by the equations:

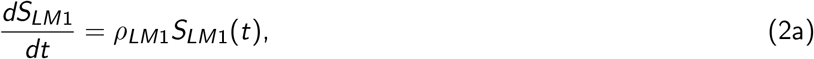

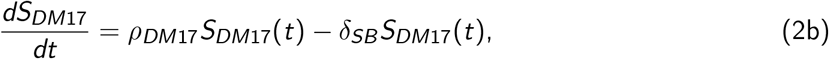

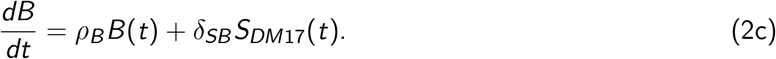

It is characterized by the set (*ρ_LM_*_1_, *ρ_DM_*_17_, *δ_SB_*, *ρ_B_*) of four parameters, following the same notation convention as in the SB model.

Finally, the SCB model (Figure 2C) also comprises the same self-renewing and differentiated compartments as the SB model, as well as a hypothetical *commited cells* compartment *C*. This compartment comprises intermediary cells that are committed to differentiation, yet not fully differentiated. The dynamics of these three compartments are given by the equations:

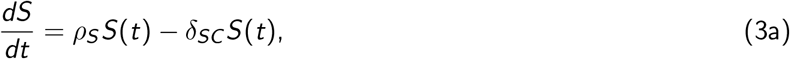

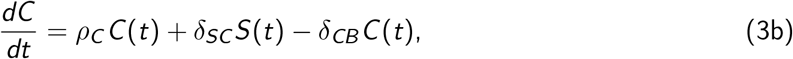

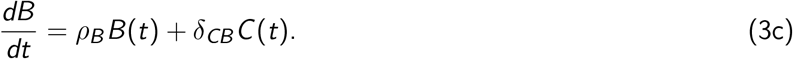

It is characterized by the set (*ρ_S_*, *δ_SC_*, *ρ_C_*, *δ_CB_*, *ρ_B_*) of five parameters, following the same naming convention as the two other models.

Moreover, it should be noted that differential systems (1) to (3) are fully linear, and that their matrices are lower-triangular, which makes them easily solvable analytically. Their simulation is thus very fast. The detail of the analytical solutions to these systems is given as supplementary material.

Finally, not all variables in the models can be measured through the experiments that we presented in section 2.1, and we only have access to two observables of the system: the total amount of living cells *T* (*t*) through trypan blue staining, and the amount of differentiated cells *B* through benzidine staining. The number of living cells *T* can be measured in LM1 and in DM17, yet in LM1 there is no differentiation, so in the LM1 experiment *T* = *S* (or *T* = *S_LM_*_1_ in the S2B model). The number of differentiated cells *B* can be measured in DM17 (it is null in LM1).

#### 2.2.2 Error Model

As will be explained below, we used the maximum likelihood approach to estimate the parameters of our model, and the profile likelihood approach to assess its identifiability. Both these methods require the proper definition of the likelihood of our model, *i.e.* the definition of a statistical model for the prediction error of our dynamical model, *y – f* (*t*, *y*_0_, *θ*), where *y* is the data and *f* the prediction from the dynamical model (which depends on time *t*, the initial condition *y*_0_ and the parameters *θ* of the model).

This prediction error is usually modelled by a gaussian distribution with a null mean^(22,38,39)^. Then, the standard deviation of the error remains to be characterized. To this aim, there are two main options^(38)^. On the one hand, one can estimate the standard deviation *σ* of the error from the data. For this estimate to be precise, one needs to have an extensive set of data, which is not our case (Figure 1C).

On the other hand, one can build a model for the standard deviation of the error. This means finding a suitable function *g*, of the time *t*, the initial condition *y*_0_, the dynamical parameters *θ* and possibly other parameters *ξ* (which we will call the error parameters), to describe this variance. The error model is then completely described as: *y_i_*,_*j*_ ⤷ 𝒩 (*f_i_* (*t_j_*, *y*_0_, *θ*), *g_i_* (*t_j_*, *y*_0_, *θ*, *ξ*)). Several simple forms have been proposed for the function *g*^(39)^ (table 1), but it is not obvious whether one should be used in general, or if the choice of *g* should be context-specific.

**Table 1:** Definition of three different error models^(39)^.

| Error model | Definition of $g$ | Error parameters |
| --- | --- | --- |
| Constant error | $\forall j \quad g_i(t_j) = a$ | $\xi = (a)$ |
| Proportional error | $\forall j \quad g_i(t_j, y_0, \theta) = b \cdot f_i(t_j, y_0, \theta)$ | $\xi = (b)$ |
| Combined error | $\forall j \quad g_i(t_j, y_0, \theta) = a + b \cdot f_i(t_j, y_0, \theta)$ | $\xi = (a, b)$ |

With this representation, the log-likelihood of the model follows as:

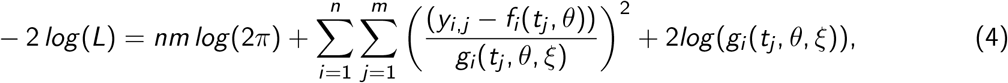

where *n* is the number of variables of the dynamic model, and *m* the number of measurement points for each variable, and from which the *log* (2*π*) term is dropped. In the end, the best-fit parameters of the model are the values of *θ* and *ξ* which minimize the quantity defined in equation (4).

### 2.3 Parameter Estimation

Considering the data at our disposal, we adopted the following procedure for parameter estimation:

1. Estimate *ρ_S_* (or *ρ_LM_*_1_ in the S2B model), and the corresponding error parameters *ξ*_1_ from the LM1 experiment. In LM1 there is no differentiation, so the *S* compartment just follows an exponential growth with rate *ρ_S_*.
2. (a) In the SB model, set *δ_SB_* so that there are no more self-renewing cells after 2 days of differentiation (which we interpret as *S*(48h) ≤ 1, or *δ_SB_* ≥ *ρ_S_* + ½*ln*(*S*(0)) from Equation (1a)). (b) In the SCB model, set *δ_SC_* so that there are no more self-renewing cells after 2 days of differentiation (which we interpret as *S*(48h) *≤* 1, or *δ_SC_ ≥ ρ_S_* + ½*ln*(*S*(0)) from Equation (3a)). This step is not an estimation step, as the value of *δ_SC_* is fixed according to the data at our disposal (Figure 1D).
3. Estimate the remaining parameters, and the corresponding error parameters *ξ*_2_, using the data from the DM17 experiment. In the SB model, the only remaining parameter is *ρ_B_*. In the S2B model, the remaining parameters are *ρ_DM_*_17_, *δ_SB_* and *ρ_B_*. In the SCB model, these are *ρ_C_*, *δ_CB_*, *ρ_B_*.

The second step of this estimation sets *δ_SC_* (*δ_SB_* in the SB model) to a value such that there are no more self-renewing cells after 2 days of differentiation. This observation does not come from the cellular kinetics experiment that we presented on Figure 1C. It rather uses the results of the commitment experiment (Figure 1D), which shows that some self-renewing cells remain in the culture after one day, but that they all are differentiated after two days^(17)^.

In the SCB model, considering that the sef-renewing compartment S is characterized by an exponential dynamic, and that there are no more self-renewing cells if and only if *S* ≤ 1, this provides an upper and a lower bound for *δ_SC_*:

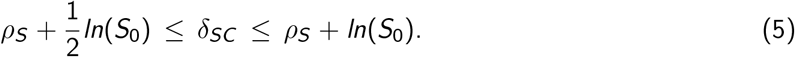

For the sake of simplicity, we will set *δ_SC_* = *ρ_S_* + ½*ln*(*S*_0_) in order to verify the equality: *S*(48h) = 1 in the following.

These considerations do not affect the S2B model, in which the switching of the culture medium only affects the proliferation rate of the S compartment.

In both estimation steps, the-log likelihood was minimized using the Truncated Newton’s algorithm^(40,41)^ implemented in the python package for scientific computing scipy^(42)^. Convergence to the global minimum was assured by a random sampling of the initial guesses for parameter values

### 2.4 Model Selection

In order to choose a proper error model, one needs to adopt a selection criterion, which allows to rank models and keep only the best ones. The most straightforward one is the log-likelihood that we presented in equation (4): the lower −2 *log* (*L*), the better the model.

Yet it only accounts for the quality of the fit of the model, without incorporating the complexity of the model. This means that –2 *log* (*L*) favours overly complex models, whereas a good selection criterion should balance the quality of the fit of the model with its complexity. This is the case for Akaike’s Information Criterion (*AIC*^(43)^):

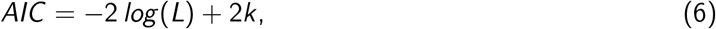

where *k* is the number of parameters of the model. Unfortunately this criterion is valid only when the sample is very large (*n* > 40*k*, where *n* is the number of points in the sample^(43)^). Corrections to *AIC* have been developed to account for the sample size, and are incorporated in the corrected *AIC*, namely *AICc*^(43)^:

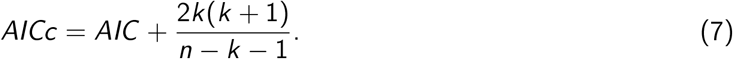

From this definition it is straightforward that *AICc* converges towards *AIC* when the sample is sufficiently large. The corrective term in *AICc* has been developed only for linear models. However, since there is no selection criterion derived from *AIC* for non-linear models, the literature recommends using *AICc* when in doubt^(43)^. As a consequence, we will use the *AICc* for our model selection, despite its statistical non-linearity (its outputs are not a linear combination of the inputs).

It is important to notice that the *AICc* value contains little information in itself, and that what matters for the purpose of model selection is the difference between the *AICc* of two competing models. Indeed, the best model in a set of models is the one with the lowest *AICc*, but if the second best model has an *AICc* very close to the minimal *AICc*, this means that this second model is close to being the best one. This is what motivates the introduction of the *AICc* difference for each model:

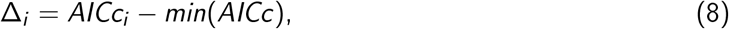

where *AICc_i_* is the *AICc* for the *i*-th model. It is admitted that a model is significantly worse than the best one if its *AICc* is more than 10 units higher than the minimal *AICc*^(43)^.

Another way of selecting the best models in a family of models is through the Akaike’s weights^(43)^:

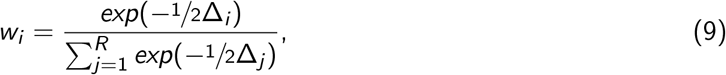

where *R* is the number of competing models. The Akaike’s weight of a given model in a given set of models can be seen as the probability that it is the best one among the set. In this setting, selecting the best models of a set of models means computing their Akaike’s weights, sorting them, and keeping only the models whose weights add up to a significance probability (for example, 95%).

We will use Akaike’s weights to select between models in the rest of this study.

### 2.5 Identifiability Analysis

The identifiability of a model describes the unicity of the correspondence between the parameters of the model and the data. In other words, a model is identifiable if and only if there is only one parameter set that reproduces a dataset, for a given criterion of the fit quality.

Though very simple, this definition is very hard to use in practice, so several methods of identifiability analysis have been developed to assess confidence intervals for their parameter estimates^(21)^. We chose to use one of these to assess the identifiability of our model, which uses the statistical notion of profile likelihood^(22,44)^.

For a model with *k* parameters, a parameter space Θ = {(*θ_i_*, *i* ∈ {1, 2, …, *N*}), *θ_i_* ∈ ℝ }, and a likelihood *L*, the **profile likelihood** *PL_θ_i* with respect to parameter *θ_i_* is defined as:

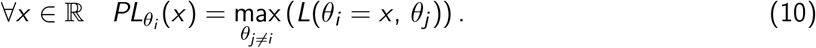

Namely, the profile likelihood with respect to a parameter at a certain value is the likelihood of the model, maximized with respect to all the other parameters. Computing the profile likelihood at a certain value *x* of a parameter *θ_i_* means to set *θ_i_* = *x* and to estimate the values of the other parameters *θ_j_* that minimize the error −2 *log* (*L*) in this setting. Consequently, −2 *log* (*PL*) is minimal at the optimal parameter values set, and increases in both directions.

In a highly dimensional parameter space where the exact shape of the likelihood landscape is not straightforward, the profile likelihood summarizes this shape along the axis of *θ_i_*, by keeping the likelihood as high as possible in all directions. It is possible to define a confidence interval *CI_θ__i_* at a level of confidence *α* ∈ [0, 1] for a parameter *θ_i_*, derived from the evaluation of the profile likelihood^(22)^:

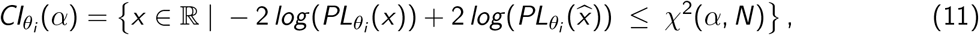

where *xˆ* is the optimal estimate of *θ_i_*, *N* is the number of parameters being estimated and *χ*^2^(*α*, *N*) is the *α*-quantile of the *χ*^2^ distribution with *N* degrees of freedom.

Namely, all the parameter sets that render a profile likelihood closer to its minimal value than a threshold *χ*^2^(*α*, *N*) belong to the confidence interval. The threshold depends both on the number of parameters estimated (the more parameters, the higher the threshold and the wider the confidence interval) and on the requested confidence level (the higher the confidence level, the higher the threshold, and the wider the confidence intervals).

Once a confidence interval has been extracted for a parameter *θ_i_*, we say that *θ_i_* is **practically identifiable at the level of confidence *α*** if and only if its confidence interval at the level *α* is bounded^(22)^.

Then we say that a model is **practically identifiable at the level of confidence *α*** if and only if all its parameters are^(22)^.

The profile likelihood approach is a good way of addressing the identifiability of a model, because it allows to detect both structural and practical non-identifiabilities. This feature makes the approach more efficient in practice than most of the other methods in the field^(21,23,45,27)^.

## 3 Results

### 3.1 Measurement error

The data that we used for the calibration of our models are displayed on Figure 1C. For readability, it displays the total cell counts in log scale, and the differentiated cell counts as a fraction of the total count. This representation emphasises the discrepancy between the cell counts in LM1 medium at the first two timepoints: the measured cell population in LM1 decreases between 0h and 24h, in both conditions (control, and rapamycin-treated).

First, every timepoint displayed on figure 1C was obtained by a single measurement. This means that they are subject to a measurement error which might blur the exponential growth that one would expect in such experiments^(35)^. Second, all the cell cultures used in this study were initiated with 25,000 cells. A sampling error in this initiation step would affect all the cell counts of the experiment.

In the course of an experiment where cells are grown in a specifically designed medium^(35)^, one would expect the cell populations to increase. We thus conclude that the observed decrease in the total living cell number in the LM1 medium (in the control and the rapamycin-treated cases) is due to the experimental error that the protocol suffers rather than to a hypothetical biological feature of the cells under study.

### 3.2 Fitting the Model with no Treatment: Model Design and Validation

#### 3.2.1 Choosing a structural model: selection approach

In order to choose the best model at reproducing the *in vitro* dynamics of erythropoiesis, we computed the maximum-likelihood estimates of the parameters of the three models we designed in section 2.2. To this end, we coupled each dynamic model (SB, S2B and SCB) with the same error model (the proportional one) and estimated their parameters (which are displayed in table S1).

For each of these three models, we computed the likelihood-based selection criteria that are displayed on Table 2. The S2B and SCB models appear as the best ones and offer very similar fits, with the SB model being far worse. It is thus possible to discard the SB model based on the Akaike’s weights, but it remains impossible, based on this criterion, to decipher which of the two remaining models should be used to best describe the *in vitro* erythropoiesis.

**Table 2:** Selection criteria evaluated for the three dynamical models presented on Figure 2. L_1_ and L_2_ are the likelihoods of the models evaluated with the data of the first and last estimation steps respectively. k is the number of estimated parameters in each of the models, according to the procedure described in section 2.3. For each model, the sample size is n = 15.

| Error model | $-2 \log(L_1)$ | $-2 \log(L_2)$ | k | AIC | AIC<sub>c</sub> | $\Delta AIC_c$ | $w_{AIC_c}$ |
| --- | --- | --- | --- | --- | --- | --- | --- |
| SB | 106 | 199 | 5 | 315 | 322.1 | 20.1 | 0 |
| S2B | 106 | 174 | 6 | 292 | 302 | 0 | 0.5 |
| SCB | 106 | 174 | 6 | 292 | 302 | 0.40 | 0.5 |

However, the S2B model does not describe the results of the commitment experiment (Figure 1D). In this model indeed, self-renewing cells switch between different self-renewing rates upon medium switching. As a consequence, switching the cells back and forth between the two media should just switch their proliferation rate, without affecting their proliferation ability. So the cells would never lose their proliferation ability, as opposed to the result of the commitment experiment.

On the other hand, the SCB model predicts that upon switching to the differentiation medium, the cells from the S compartment start differentiating. Once they are all differentiated, cells from the C and B compartments can still proliferate, but this proliferation might be cancelled by a switch back to the LM1 medium. It is thus impossible to describe the process of commmitment with the S2B model, while it is possible with the SCB model.

As a conclusion, the SCB model is the best-fitting model which also accounts for the results of the commitment experiment, making it our dynamic model of choice for the rest of this study.

#### 3.2.2 Choosing an error model: selection approach

Following the same approach for selecting the error model, we computed the best-fit estimates of the parameters of the three error models presented in Table 1, coupled with the SCB model. These parameter values are given in Table S2, and the values of the corresponding selection criteria are given in Table 3.

**Table 3:** Selection criteria evaluated for the three error models presented in table 1. L_1_ and L_2_ are the likelihoods of the models evaluated with the data of the first and last estimation steps respectively. k is the number of estimated parameters in each of the models, according to the procedure described in section 2.3. For each model, the sample size is n = 15.

| Error model | $-2 \log(L_1)$ | $-2 \log(L_2)$ | k | AIC | AIC<sub>c</sub> | $\Delta AIC_c$ | $w_{AIC_c}$ |
| --- | --- | --- | --- | --- | --- | --- | --- |
| Constant Error | 107 | 195 | 6 | 314 | 325 | 22 | $1.48 \times 10^{-5}$ |
| Proportional Error | 106 | 174 | 6 | 292 | 302 | 0 | 0.99 |
| Combined Error | 106 | 174 | 8 | 296 | 320 | 18 | $1.58 \times 10^{-4}$ |

The proportional error model appears as the best one, since it offers the best fit to the data, with the fewest parameters.

As a consequence, in the rest of this paper, we will consider the SCB model with proportional error.

#### 3.2.3 Identifiability analysis

In order to use a model for predictive purposes, one needs to assess its identifiability. The profile likelihood curves of all estimated parameters are displayed on Figure 3, for the SCB model with proportional error. For *ρ_S_* and *b*_1_, which are estimated together in the first step of the procedure, the identifiability threshold at confidence *α* = 0.95 is *χ*^2^(0.95, 2) = 5.99. For *ρ_C_*, *δ_CB_*, *ρ_B_* and *b*_2_, which are estimated together at the last step of the estimation procedure, the threshold is *χ*^2^(0.95, 4) = 9.49.

**Figure 3:**
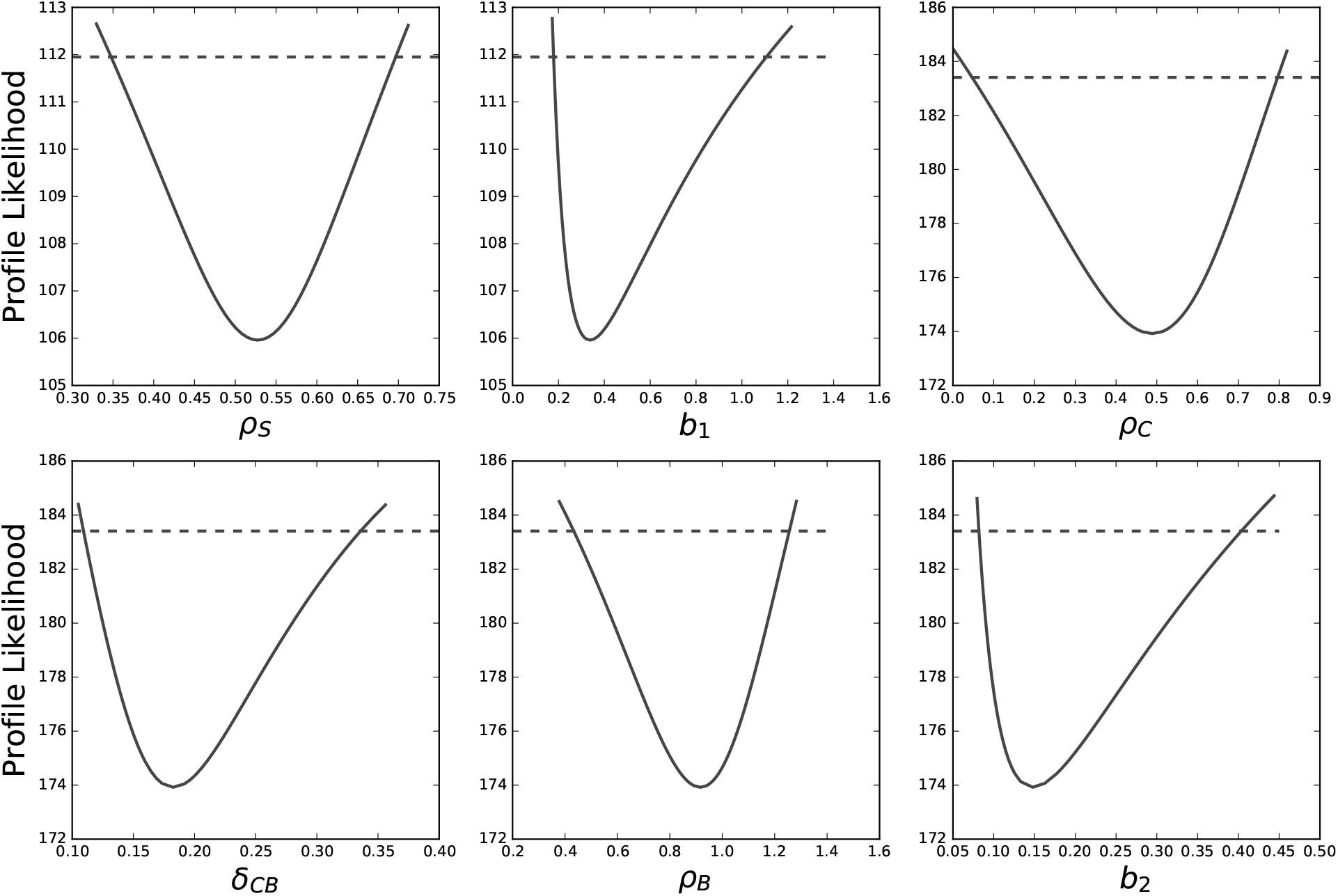
The SCB model with proportional error is fully identifiable. Solid curves are the profile likelihood curves of each estimated parameter of the model. Dashed lines give the identifiability threshold of each parameter at confidence level *α* = 0.95.

For every parameter of the model, the profile likelihood curve crosses the threshold on both sides of the optimum, which means that every parameter of the model is identifiable, at the level of confidence *α* = 0.95. The confidence intervals of the parameters^(22)^, extracted from these profiles (equation (11)) are displayed in Table 4.

**Table 4:** Confidence Intervals of the parameters of the SCB model with proportional error. Displayed are the confidence interval boundaries at level *α* = 0.95, extracted from figure 3, and the best-fit estimate of all the estimated parameters of the model. For *δ_SC_*, which is not estimated, no optimal value can be computed, but absolute bounds on its values can be computed with equation (5). Parameters are grouped by their estimation step in our procedure: *ρ_S_* and *b*_1_ are estimated together in the first step, then *δ_SC_* is set, and finally the four other parameters are estimated together. For the proliferation rates *ρ_S_*, *ρ_C_* and *ρ_B_*, we also give the corresponding doubling times of the populations (*i.e.* how long would it take to double the population in the absence of differentiation?). For the differentiation rates *δ_SC_* and *δ_CB_*, we also give the half-life of the corresponding populations (*i.e.* how long would it take to differentiate half the cells from the undifferentiated population, in the absence of proliferation?)

| Parameter | Lower bound | Time (h) | Optimal value | Time (h) | Upper bound | Time (h) |
| --- | --- | --- | --- | --- | --- | --- |
| $\rho_S$ | 0.35 | 48 | 0.53 | 31 | 0.70 | 24 |
| $b_1$ | 0.18 | - | 0.34 | - | 1.1 | - |
| $\delta_{SC}$ | 5.6 | 3 | - | - | 11 | 2 |
| $\rho_C$ | 0.049 | 340 | 0.49 | 31 | 0.80 | 21 |
| $\delta_{CB}$ | 0.11 | 150 | 0.18 | 92 | 0.34 | 49 |
| $\rho_B$ | 0.44 | 38 | 0.92 | 18 | 1.3 | 13 |
| $b_2$ | 0.081 | - | 0.15 | - | 0.41 | - |

For a given parameter, the size of the confidence interval depends on the number of parameters that are estimated together, the required level of confidence, and the likelihood function used for the computation. By definition (equation (10)), the profile likelihood renders as large of a confidence interval as possible, because it increases as slowly as possible on each side of the optimal parameter value. This means that identifiability is harder to satisfy with the profile likelihood approach than with other definitions, for example based on a linearization of the likelihood surface at the optimal parameter set^(22)^.

As a consequence, the fact that the parameter confidence intervals presented in Table 4 may appear as quite large is not a sign that the parameters are poorly estimated. It is rather the evidence that they remain identifiable even with a very stringent definition of identifiability, and at a high level of confidence.

It is not possible to study the identifiability of *δ_SC_*, since it is not estimated from the data. However, the commitment experiment (Figure 1D) give us access to a lower and an upper bounds for its value (equation 5). We determined the optimal likelihood of the model, and its optimal parameters in this range of value (figures S1 and S2), showing that the choice of *δ_SC_* does not influence the dynamics of our model.

### 3.3 Modelling differentiation in the control case

A simulation of the model with the identified values of its parameters, is reproduced, with the corresponding experimental data, on Figure 4A. The overall quality of the fit is good, especially for the DM17 populations. Regarding the self-renewing cells in LM1 medium though, the fit seems more qualitative than quantitative, due to technical imprecision of the measure (see section 3.1).

**Figure 4:**
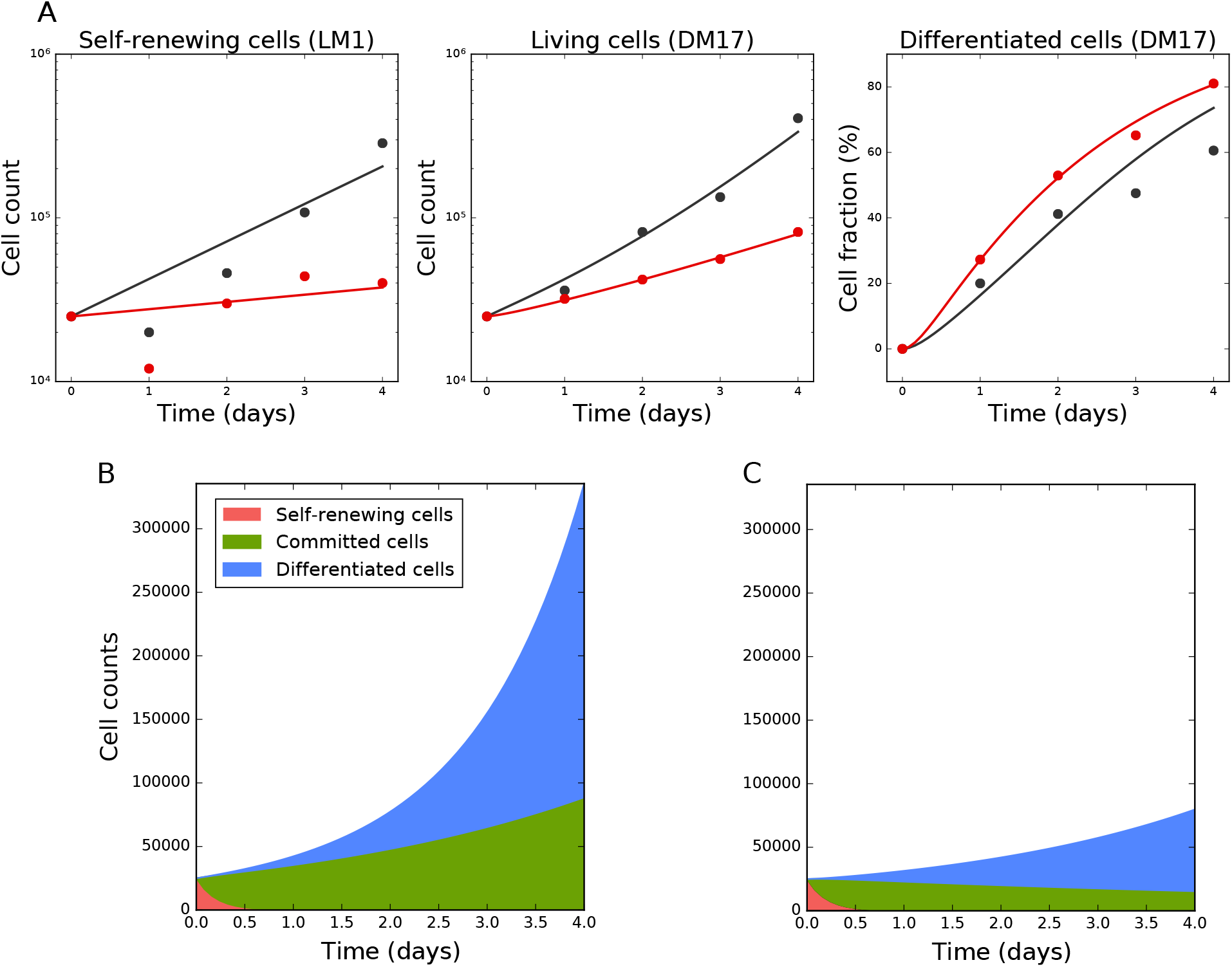
The model reproduces the cellular kinetics observed *in vitro*. A: Simulation of the SCB model with proportional error in the untreated (black) and rapamycin-treated cases (red). Solid lines represent a simulation of the SCB model with proportional error, with its best-fit parameters. Dots are the experimental data. Displayed are the total number of living cells in LM1 and DM17 media (in log-scale), and the fraction of differentiated cells in DM17. B-C: Numbers of cells in each compartment as a function of time in the untreated (B) and treated (C) cases.

The precise values of each parameter of the model are reported in Table 4. The proliferation rate of the committed cells is slightly lower than the one of the self-renewing cells (the net doubling-time of the committed compartment is about 34h, whereas the doubling-time of the self-renewing compartment is around 31h). The differentiated cells proliferate with a higher rate (their doubling time is about 18h). Taken together, these doubling times explain the faster proliferation of cells in the DM17 medium than in the LM1, in agreement with previous data^(35)^. Finally, the self-renewing cell differentiation is very fast: the lowest possible value of *δ_SC_* gives them a half-life of 3h in DM17. This means that half of the S compartment would differentiate every 3h in the absence of proliferation. On the contrary, the differentiation of the committed cells is much slower (if they stopped proliferating, they would have a half-life of 90h).

The timescales at which these processes occur are pictured on Figure 4B, which displays the number of cells in each state during a simulation of the model. As specified by the setting of the value of *δ_SC_*, the population of self-renewing cells quickly collapses, and the culture becomes a mixing of committed and differentiated cells. Both of these compartments then grow at their own rate.

At this stage, we have developed a very simple model of erythroid differentiation, that accounts well for the data used to calibrate it. Plus, it is fully identifiable (at confidence level 95%). We thus use it to study the effect of rapamycin, a drug known to affect the *in vitro* erythroid differentiation^(36,37)^.

### 3.4 Modelling differentiation under Rapamycin treatment

Rapamycin is known to increase the proportion of differentiated cells in cultures of chicken erythroid progenitors^(36,37)^ (Figure 1C). Yet this effect might have several origins: a decreased mortality of the differentiated cells, or an increased differentiation rate for example. To decipher between these different possible effects of the rapamycin treatment, we estimated the values of the parameters of our model in the rapamycin treated case.

To avoid an overparameterization of the rapamycin effect, we considered that for each estimated parameter of the model, the value under rapamycin treatment could be either equal to the value in the untreated case, or equal to another value yet to be estimated. The first option would not introduce a new parameter in the model, but the latter would. Our model has seven parameters (5 dynamical parameters and 2 error parameters), of which 6 only are estimated (since *δ_SC_* is entirely determined by the value of *ρ_S_*). This means that we can define 2^6^ = 64 models of rapamycin treatment, by keeping some of the parameters unchanged compared to the untreated case, and re-estimating the others with the data presented on Figure 1C.

We thus estimated the parameters of these 64 possible models of the treatment, and computed their likelihood-based selection criteria, the same way as we did for the dynamic model and the error model (see section 3.2). The Akaike’s weights of the best three of these models are shown on Figure 5A. The best model of the treatment is responsible for 94 % of the weight of the 64 models, making it by far the best model for the rapamycin treatment. A simulation of this model is displayed on Figure 4, which indeed shows the quality of its fit to the data obtained under rapamycin treatment.

**Figure 5:**
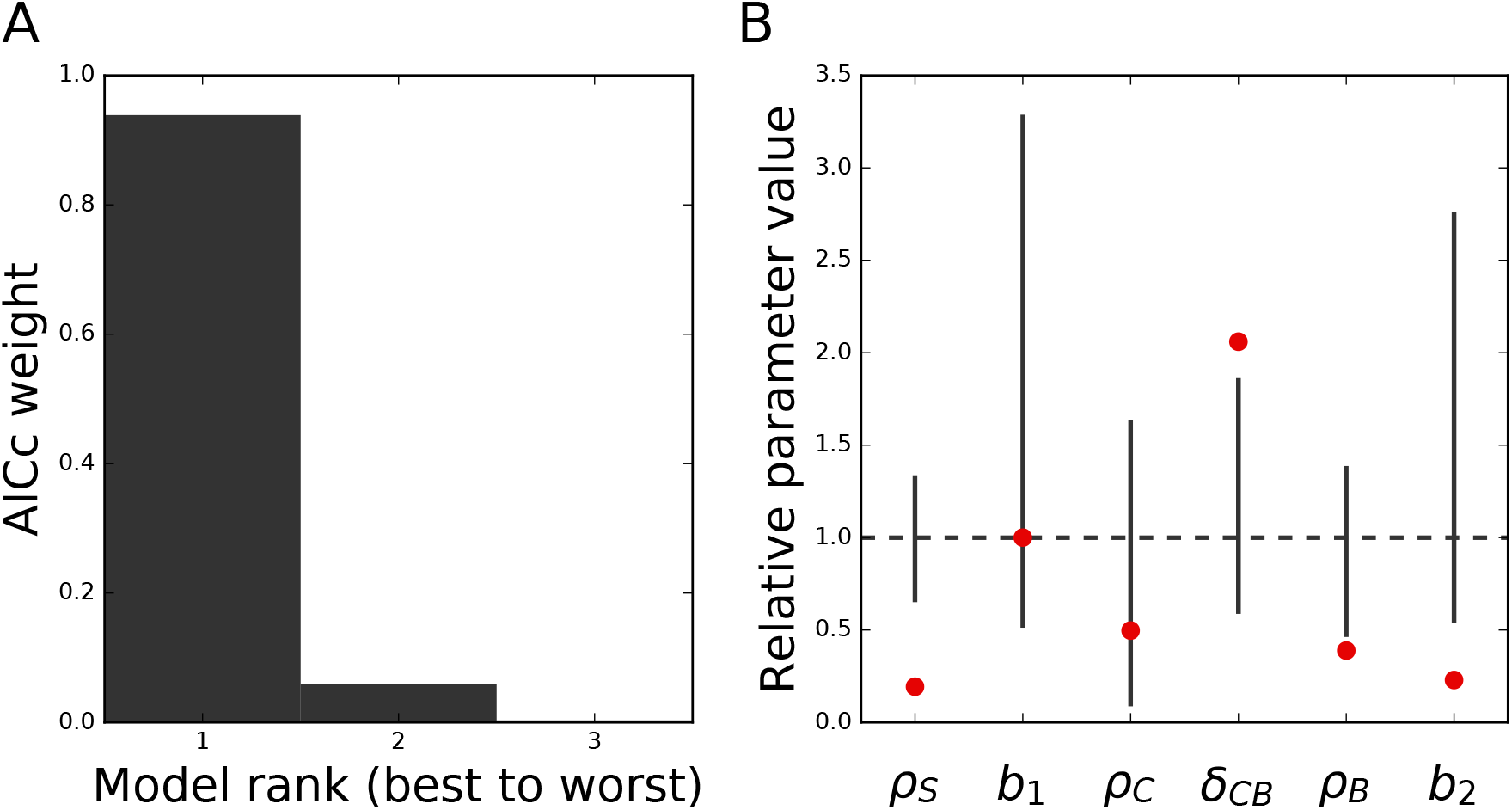
Modelling erythropoiesis under rapamycin treatment. A. Akaike’s weights of the three best models of the rapamycin treatment. The 61 other models are not displayed for readability. B. Parameter values in the best model of rapamycin treatment. Red dots are the ratio of the parameter values under rapamycin treatment with their values in the untreated case. Straight lines represent the confidence intervals of the values in the untreated case, computed from Figure 3. The dashed line indicates the parameter values in the untreated case, by which all parameters are scaled for readability.

This model is obtained by varying all the parameter values except *b*_1_, compared to the control case, as displayed on Figure 5B. Moreover, 4 of the other parameters are varied outside of their 95% confidence intervals, showing the strength of the treatment effects. The three proliferation rates *ρ_S_*, *ρ_C_* and *ρ_B_* are reduced under the treatment, while the differentiation rate *δ_CB_* is increased.

Finally, the effect of rapamycin on the repartition of cells between the different compartments is displayed on Figure 4C. Under rapamycin, the C compartment decays, when it proliferated in the control case. Moreover the B compartment has a longer doubling time under rapamycin treatment (48h instead of 18h in the untreated case). These two effects explain why the drug treatment reduces the overall amount of cells and increases the proportion of differentiated cells in the culture.

## 4 Discussion

We proposed a model for the *in vitro* erythroid differentiation, which comprises two components. First, the dynamic component is the set of ODE written in equation (3), which describes the dynamics of three cell populations. Second, we added an error component which describes the distribution of the residuals of the dynamic model,

The three populations of our dynamical model are related to three different stages of differentiation of the progenitors. The first one is in a self-renewal state S where differentiation has not started, and the third one has finished differentiating. The second population lies in the middle, in a state of commitment C where cells are not fully differentiated yet, but cannot go back to self-renewal.

Similar 3-states models have already been used to describe differentiation^(19)^. Their success probably stems from the fact that it would be difficult to describe differentiation as the transition between only 2 states, as we highlighted in the context of *in vitro* erythropoiesis with our SB model. Actually, differentiation from one cell type to another is a continuous process, so its best description would probably be a continuum of states, which would punctuate the transition between the two cell types^(19,46)^.

However, explicitly accounting for this continuum would require an infinity of intermediary states. For example, the levels of differentiation factors inside the cell could be used as a measure of its differentiation state^(47,48)^. Such kind of a model should be able to describe differentiation more faithfully than ours. Yet our model, though simplistic, reproduces our experimental data quite well, and is identifiable. Moreover, since it is fully linear and thus analytically integrable, its simulation and calibration are very quick.

Such simplicity and identifiability of our model would probably make it valuable to describe differentiation in other contexts.

Once the model was chosen, we verified the accuracy of its parameter estimates. We showed that among the seven parameters of our model, the six that are estimated by the maximum-likelihood approach are identifiable, and that the choice of the seventh one does not alter the behaviour of the others. Using the Profile Likelihood approach, we computed confidence intervals for our parameters. Even though their relatively large size might be interpreted as a lack of accuracy in the estimates of the parameters, it is not the case since the identifiability of a parameter is harder to satisfy by the Profile-Likelihood approach than using other methods. We thus showed that our model is fully identifiable, even using a very stringent criterion.

After demonstrating the validity of our model in the control case, we used it to study the effect of rapamycin, a chemical drug which is known to impact the differentiation of erythroid progenitors^(36,37)^. We designed 64 different models of the rapamycin treatment, which differ by the combinations of parameters that are affected by the treatment. Evaluating the quality of their fit to the data allowed us to retain only the best model of rapamycin treatment. The parameter values in this model reveal that rapamycin increases the differentiation rate of the intermediary cell compartment, and reduces the net growth rates of the three other compartments. This means that rapamycin increases the differentiation of the cells in culture, and also affects the balance between their proliferation and mortality. The reduced net proliferation rates might be caused by a reduced proliferation, an increased mortality or any joint variation of the two processes equivalent to one of these effects (e.g. reduced mortality, and an even more reduced proliferation).

In the context of other perturbations of differentiation (for instance, treating cultures with a different drug than rapamycin), should the drug influence be less strong, we might need a more subtle means of parameter evaluation, such as the fused lasso penalized regression^(49)^.

At the moment, our approach suffers one major drawback that is the size of our dataset. Indeed, the precision of our prediction of cell numbers at one timepoint relies on the precision of our measures of these numbers. And this measurement precision is directly related to the number of repetitions of the measurements which are performed by the experimentalist.

Repeating the same experiment several times would thus increase measurement precision and average out measurement noise. This would allow a more precise estimation of the error model parameters, and in turn would increase the precision of the dynamic model. What we call *repeating the same experiment* here does not simply consist in counting cells from the same time point several times to average out sampling biases. It rather involves putting new cells in culture and following their populations over time, as two full replicates of the experiment.

In this setting, the measurement error would not just be limited to technical noise due to the sampling of cells from the culture for counting. It would rather be related to differences between the kinetic features of the cells in culture, *i.e.* to actual biological heterogeneity. This heterogeneity would be averaged by the estimation of one parameter set to fit all the data.

One way of accounting for this heterogeneity in a model, without averaging it out during parameter estimation, is through the use of mixed effect models, that is a mathematical model (e.g. a set of deterministic ODE, like ours) whose parameters are modeled by distributions of random variables^(39)^. We are presently assessing the ability of such mixed effect models to characterize both the behaviour of the cells in culture on average, as well as their variability.

## 5 Acknowledgements

We thank members of the SBDM and Dracula teams, as well as Dr. Boudaoud (ENS Lyon), Dr. Leclercq Samson (Laboratoire Jean Kuntzmann, Grenoble, France) and Dr. Raue (Merrimack Pharmaceuticals, Cambridge, Massachussetts) for enlightening discussions.

We also thank Pr. Saccomani (University of Padova, Italy) for providing the manuscript for reference^(20)^.

Finally, we want to thank the BioSyL Federation and the LabEx Ecofect (ANR-11-LABX-0048) of the University of Lyon for inspiring scientific events.

## 6 Supplementary Materials

The supplementary materials file shows the analytical solutions of our three dynamic models, the parameter values estimated for the three dynamic models (table S1) and the three error models (table S2), as well as the influence of our choice of *δ_SC_* on the likelihood of the model (figure S1) and on its parameter values (figure S2).

All the datasets and pieces of code analysed and generated during the current study are available in a public github repository, at https://github.com/rduchesn/Dynamic_Model_Erythropoiesis.

